# Pangenomic exploration of *Theobroma cacao*: New Insights into Gene Content Diversity and Selection During Domestication

**DOI:** 10.1101/2023.11.03.565324

**Authors:** Xavier Argout, Gaetan Droc, Olivier Fouet, Mathieu Rouard, Karine Labadie, Bénédicte Rhoné, Gaston Rey Loor, Claire Lanaud

## Abstract

The Cacao tree (*Theobroma cacao L.)* is a major cash crop and an important source of income for millions of farmers across Africa, Asia and Latin America. However, cacao farmers and producing countries are facing multiple challenges including pest and disease management, sustainable production under threat of climate changes and diversification of cocoa flavor profiles. Addressing these challenging requires a deeper understanding of the existing genetic diversity of the species. Yet, very little is known about the intraspecific gene content variation.

In this study, we used the genome of 216 accessions of *T. cacao* (including 185 newly re-sequenced) covering a broad genetic diversity of the species to construct the first pan-genome of the cacao tree. As a result, we predicted a total of 30,489 pan-genes, enriching the reference Criollo genome with 1,407 new genes.

Our analysis revealed that only a small fraction of these genes (9.2%) is dispensable, being absent in some individuals. However, these genes could represent a valuable resource for breeding efforts aimed at improving disease resistance in the species.

We used this new pangenome resource to gain insights into how diversification and domestication processes have influenced genomic variability within the species. Notably, we observed a significant loss of genes within the domesticated Criollo genetic group. Furthermore, we found evidences suggesting that domestication processes have had an impact on the vigor and disease tolerance of Criollo accessions. In summary, our research has contributed to a better understanding of the cacao tree’s genome diversity. These findings offer new avenues for biological discovery and breeding, ultimately addressing the challenges faced by cacao farmers and producing countries.

## Introduction

*Theobroma cacao L.*, commonly known as the cocoa tree, is a perennial diploid species native to the upper Amazon basin and is a pivotal cash crop for millions of smallholder farmers in countries across the humid tropical regions of the world^1^. The cocoa sector is currently facing many challenges which include low productivity^2^, significant production losses due to pests and diseases^3^, contamination by heavy metals such as Cadmium^4^, impacts of climate change^5^ and deforestation concerns^6^. Furthermore, in recent years, consumer preferences have shifted towards high-quality chocolate products using sustainable practices. This trend has prompted farmers and manufacturers to search for diverse and unique cocoa flavor profiles, and to produce fine and flavored cacao beans. Comprehensive exploration of the *T. cacao* genetic diversity at genomic level provides resources that can help in addressing these challenges. Effectively utilizing this information should support breeding programs to develop varieties adapted to these different environmental contexts^7^.

Over the past decade, significant progresses have been made on cocoa genetic improvement through the generation and utilization of genomic data. To date, annotated chromosome-scale genomes of two *T. cacao* accessions are available: the B97-61/B2 Criollo reference genome produced in 2011^8^, further improved in 2017^9^ and the genome of Matina1-6, an Amelonado cultivar sequenced in 2013^10^. These two genomes have significantly advanced scientific discoveries in *Theobroma cacao*, allowing for instance the development of molecular markers and the candidate genes identification through GWAS or gene expression analysis using NGS^11^. They also enabled substantial progress in modern cocoa breeding approaches, using marker assisted or genomic selection. Nevertheless, recent studies in plant genomics indicate that one or few reference genomes cannot capture the full range of genomic diversity of plant species, including the presence/absence of protein coding genes sequences, which may lead to a bias associated with the use of a single reference^12^. The development of the pan-genome, defined here as a representation of all DNA sequences within a species, can address this issue.

Plant pan-genomes have been increasing dramatically in recent years^13^. The concept of pan-genome includes core genes, found in all individuals, as well as variable genes (or dispensable genes), which are absent in some individuals or genetic groups. Gene presence and absence variations (PAVs) are an important source of genetic divergence at a population level and their understanding can support applications for crop improvement because they can contribute to trait variation. Functional analysis of variable genes in multiple crops have revealed their enrichment in genes related to disease resistance, abiotic stresses, environmental adaptation, and quality when compared to gene core fraction^14–17^.

Very little is known about the complete genomic content across *T. cacao* species. In comparison to other plant species, the cocoa tree genome size is relatively small (430Mb) but can vary by about 10% depending on the genotype^8^. The genetic diversity of the species has been widely studied, particularly using microsatellites and SNPs as molecular markers^18–21^. The current genetic diversity of the species includes several genetic groups that could have diversified through isolation in refugia during the last glaciation followed by human selection and domestication^18,22^. Among the different genetic groups observed, the Criollo, highly regarded by the global cocoa industry, stands out due to its nearly unique and highly homozygous genotype, resulting from domestication processes along with genetic drift after transport from South to Central America^23^.

Recently, analyses of the genetic diversity among native cocoa trees collected during surveys in the Ecuadorian Amazon have broadened our understanding of the diversity of the species and improved the knowledge of the global genetic structure of *Theobroma cacao*. In this work, we utilized a subset of these genetic resources, along with representative samples from all other genetic groups, to construct the first *Theobroma cacao* pan-genome. This pan-genome was based on deep whole genome resequencing data generated from 216 accessions, encompassing the known diversity of the species up to the present day. We comprehensively investigated gene PAVs inside the diversity of the species and captured new protein coding genes not included in the reference genomes. We also examined how the processes of diversification and domestication of the *Theobroma cacao* genetic groups have shaped their gene content and searched for genes under selection during the domestication process of the Criollo group. Through this work, we aim to provide insights and tools that can guide future efforts in cocoa breeding and conservation strategies.

## Results

### De novo assembly of *Theobroma cacao* accessions and pan-genome construction

We selected a total of 216 *T. cacao* accessions (Figure 1a, Supplementary Table 2) covering a broad genetic diversity of the species^21,24,25^. Among these accessions, we newly sequenced the genome of 185 cocoa accessions by Illumina technology (2x150bp reads) that we combined with Illumina genome sequences of 31 accessions retrieved from public databases (Bioproject PRJNA558793). After low quality and sequencing adapters removal, read coverage per accession ranged from 40x to 75x with a mean of 58x.

**Figure 1.**
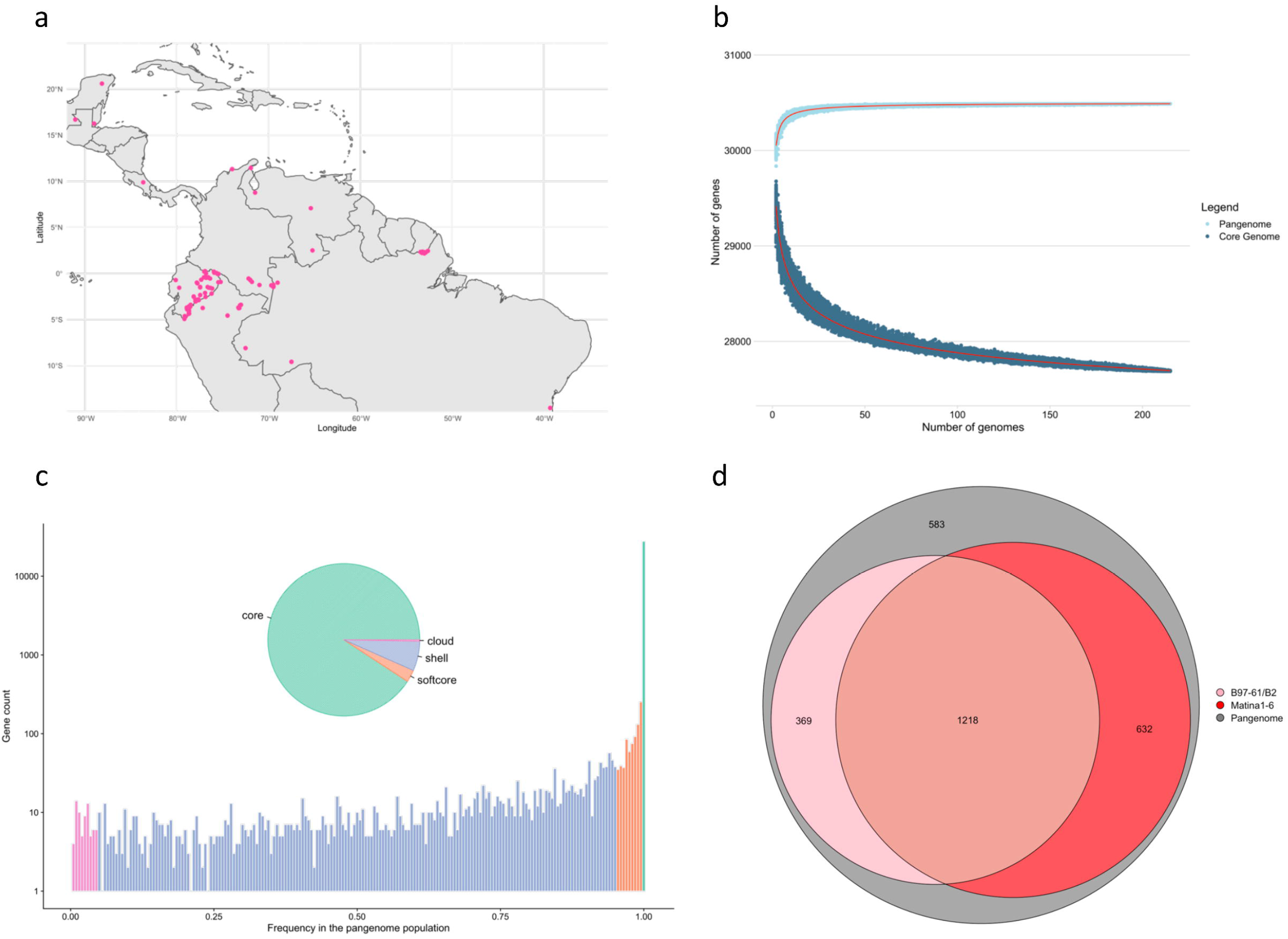
Pan-genome of Theobroma cacao. (a) Geographical distribution of the 216 T. cacao accessions used for pan-genome construction. (b) Pan-genome saturation curve indicating the distribution of the number of pan and core genes along with different numbers of sequenced individuals. (b) Pan-genome gene classification. (d) Comparison of variable genes detected in the T. cacao pan-genome with the Criollo (pink) and Amelonado (red) annotated T. cacao genomes.

We performed *de novo* assembly for each of the 216 accessions and generated a total of 76.24 Gbp of contig sequences longer than 500 bp with a N50 value of 13,357 bp. The size of the assembly ranged from 309.0 Mbp to 505.5 Mbp per accession with a mean of 354.6 Mbp (Supplementary Table 2). All accessions showed assembly completeness assessed by BUSCO higher than 92.4%, with a mean for all *de novo* assemblies of 95.2%, a value similar to that reported for the B97-61/B2 reference genome^9^ (97.3%).

The process of identification of sequences not included in the B97-61/B2 reference genome^9^ resulted in a total of 1.59 Gbp of cleaned non-reference sequences. The final non reference genome dataset was consolidated as 48.2 Mbp of sequences longer than 1,000 bp after redundancy removal.

A total of 1,407 candidate protein-coding gene models were predicted in the newly assembled contigs. From them, 99.2% exhibited significant similarities (evalue > 1e-5) with proteins of the NR database^26^ and 93.2% were annotated with gene ontology terms. Gene ontology analysis revealed that the largest groups of genes included in the non-reference genome were involved in defense response, hydrolysis of proteins, protein phosphorylation, intracellular signal transduction and production of siRNA (Supplementary Figure 1).

The final *Theobroma cacao* pan-genome, including the B97-61/B2 reference and non-reference genome dataset, had a total size of 372.9 Mbp and contained 30,489 protein-coding gene models. The pan-genome saturation curve (Figure 1b) shows that the total number of pan-genes increases as additional genomes are added, until a plateau is reached at about 50 accessions. It indicates that our sampling led to a closed pan-genome and that the diversity of *Theobroma cacao* was largely captured by the selected set of accessions. We also observed a continuously decrease of the core-genome size with the increase of population size, until a substantial decrease of the slope of the curve detected from 100 accessions, suggesting that the large number of accessions included in our dataset was large enough for an accurate characterization of the core-genome.

### Core and dispensable genome

We then categorized the presence and absence of each protein-coding gene model according to their presence frequencies among the 216 *T. cacao* accessions (Figure 1c). A vast majority of genes, 27,687 (90,8%), were defined as core genes while 2,802 were identified as variable. Variable genes were composed of 806 (2,6%) softcore genes, absent in at least one accession but present in more than 95% of the accessions, 1,924 shell genes (6,3%) present in 5-95% of the accessions, and 72 cloud genes (0,3%) present in less than 5% of the accessions. Interestingly, none of the genes were specific to a single accession, suggesting the high level of representativeness of the *T. cacao* pan-genome provided by the 216 accessions.

The comparison of variable genes detected in this pan-genome analysis with the 2 annotated genomes^9,10^ indicated that the pan-genome comprised 583 *Theobroma cacao* protein coding gene models not previously cataloged (Figure 1d). We also observed that a substantial part of the variable genes is absent in either Matina1-6 (34%) or B97-61/B2 (43%) genomes, highlighting the relevance of this new resource for the understanding of the genomic basis of traits in the *Theobroma cacao* species.

### Gene PAVs population genetic structure

We first examined the genetic structure of the population with the protein-coding gene presence and absence variations (PAVs) matrix build from the 2,802 variable genes found in the 216 *T. cacao* accessions. From admixture cross entropy criterion, we identified a minimum at K=15 (Figure 2a) indicating that the studied population comprised 15 genetic clusters. Out of these 15 genetically differentiated groups, 11 were identified and consistently named in accordance with Motamayor *et al.,* 2008^18^ and Fouet at al., 2022^21^: Amelonado, Caqueta, Contamana, Criollo, Curaray, Guiana, Iquitos, Marañon, Nacional, Nanay and Purús. The remaining four genetic clusters newly identified were named based on their respective geographical locations:, Apaporis, Pangui, Napo and Tiwinza.

**Figure 2.**
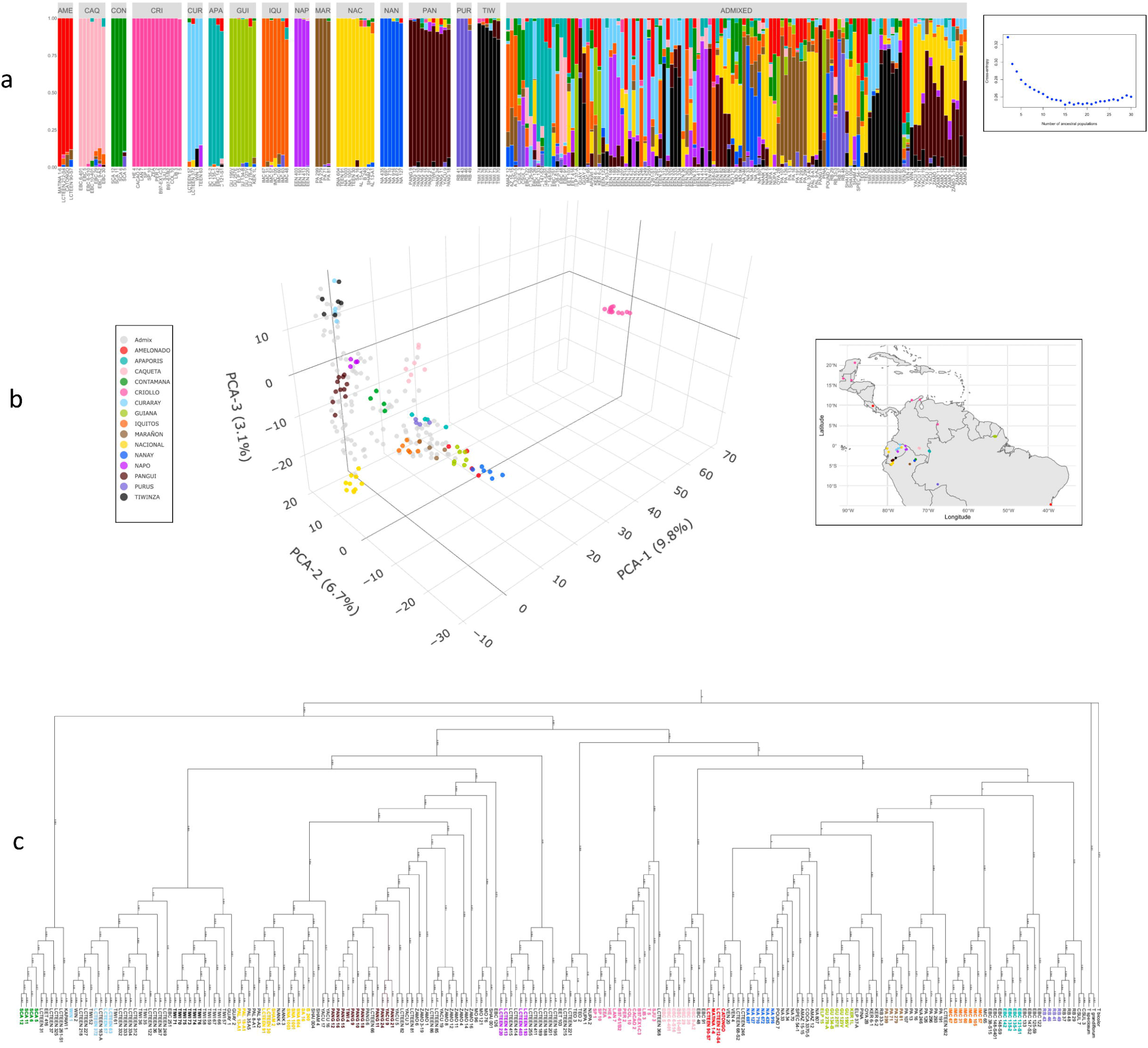
Population structure and phylogeny of the 216 T. cacao accessions based on gene PAVs. (a) Model-based clustering analysis with ancestral kinship (K = 15). The y-axis quantifies cluster membership and the x-axis lists the different accessions. Cluster AME=Amelonado, CAQ=Caqueta, CRI=Criollo, CUR=Curaray, APA=Apaporis, GUI=Guiana, IQU=Iquitos, NAP=Napo, MAR=Marañon, NAC=Nacional, NAN=Nanay, PAN=Purus, TIW=Tiwinza. (b) 3D-PCA scatter plot of the first three principal components. (c) Neighbor-joining tree was calculated from PAV distance matrix computed using the Sokal & Michener algorithm. Numbers indicate branch length and colors genetic group. The phylogenetic tree was edited using iTOL v6 (https://itol.embl.de)

Notably, Caquetá and Pangui genetic groups dissociated early from other accessions at a low K value (respectively K=4 and K=8) (Supplementary Figure 2), suggesting that these genetic groups comprise uncommon gene space among the genetic diversity of *Theobroma cacao*. Conversely, the accessions belonging to the Tiwinza genetic group dissociated from the Curaray genetic group only at K=15.

We then estimated the individual admixture coefficients and identified 95 accessions exhibiting ancestry higher than 85% to their respective identified genetic cluster (Figure 2a). These accessions were retained to investigate the genomic specificity of the genetic groups while the remaining 121 accessions with a coefficient of membership lower than 85% were considered as admixed.

The first axis of the PCA analysis (Figure 2b) clearly differentiated the domesticated Criollo group from the other genetic groups while the second axis separated roughly the genetic groups from East to West according to their geographical coordinates. Furthermore, the third component of the PCA analysis revealed a distinct pattern of separation among genetic groups based on their South to North geographic coordinates, notably pronounced in the accessions collected in Ecuador. The PCA analysis also indicated high similarity between the Curaray and Tiwinza genetic groups as the accessions from these 2 genetic groups overlap in the projection of the 3 main principal axes.

Finally, phylogenetic analysis of the 216 *T. cacao* accessions was consistent with admixture and PCA analyses (Figure 2c). The Neighbor-joining tree supported 15 clades, grouping accessions similarly to the 15 groups predicted by the admixture analysis.

### Selection of gene PAVs during *T. cacao* diversification

We further investigated whether the processes of diversification of the *Theobroma cacao* genetic groups have shaped their gene content by exanimating the distribution of gene PAVs into the 15 groups. We found that accessions of the Iquitos and Napo genetic groups had the highest gene content (gene mean number of 29835 and 29806 respectively) (Figure 3A), suggesting that the differentiation processes leading to these two genetic groups had the least impact on their gene content. At the opposite, the domesticated Criollo genetic group encoded significantly (Newman-Keuls test at p<0.05) less genes than other genetic groups (29298), suggesting that the domestication process of *T. Cacao* has led to gene loss.

**Figure 3.**
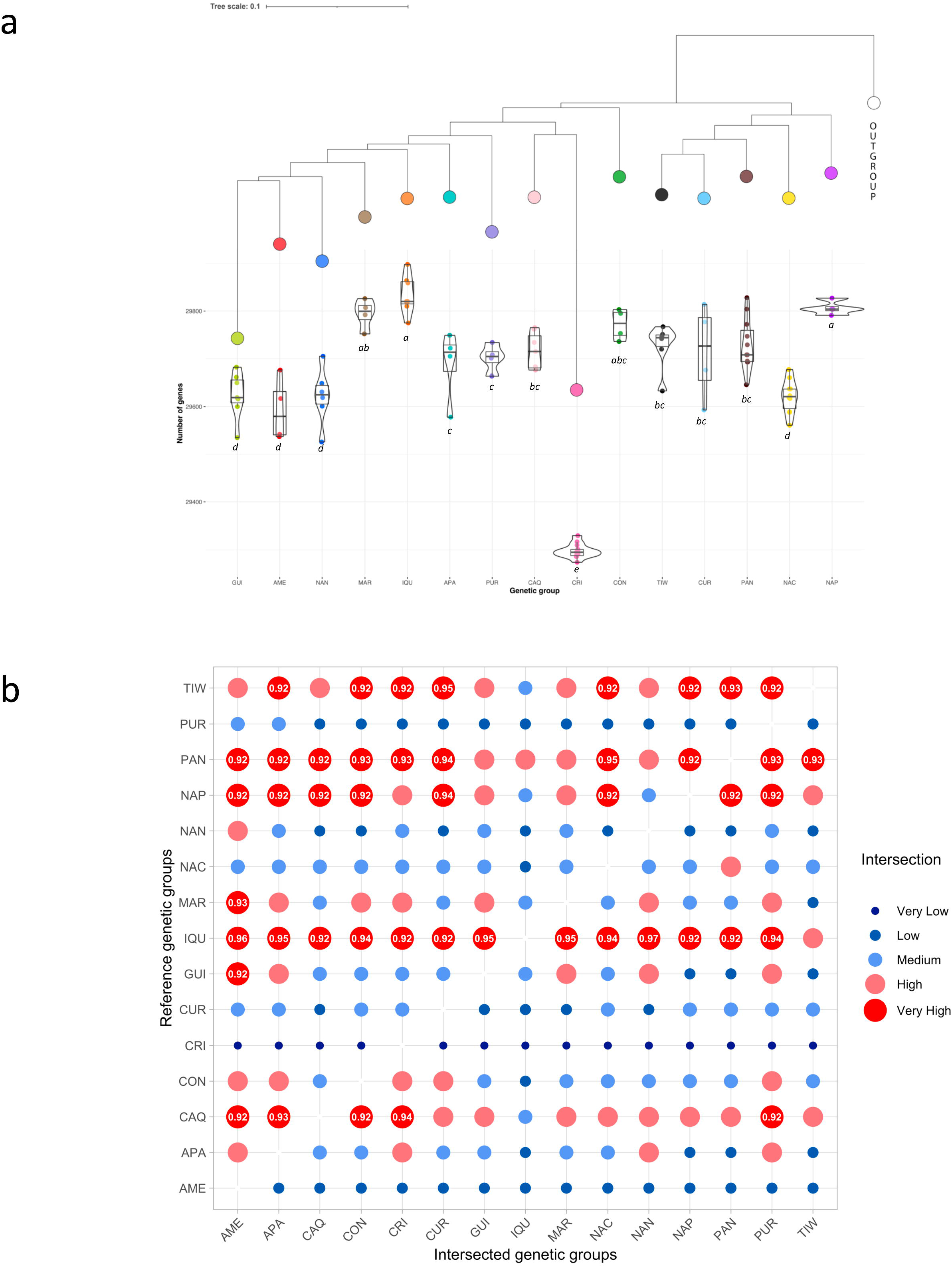
Gene number and gene space intersection among the T. cacao genetic groups. (a) Violin plots showing gene abundance for the 95 accessions exhibiting ancestry higher than 85% to their respective identified genetic cluster. Neighbor-joining tree was calculated from PAV distance matrix between the 95 accessions. Letters indicate comparable means according to the test of Newman-Keuls (probability p = 0.05). (b) Intersection plot of the gene space of the 15 genetic groups. “Very Low” indicates intersection <0.7, “Low” >=0.7 and <=1^st^ quartile (0.85), “Medium” >1^st^ quartile and <=2^nd^ quartile (0.88), “High” >2^nd^ quartile and <=3^rd^ quartile (0.91), “Very High” > 3^rd^ quartile.

Since some admixture was observed in several accessions of the 15 genetic groups (Figure 2a), we assigned a gene to a genetic group if it occurred in at least 25% of its comprised accessions. We then compared the intersection between the representative gene space of the 15 genetic groups (Figure 3b). We observed that very few genes were specific to a genetic group (Supplementary Figure 3), highlighting the fact that the boundaries between genetic groups are porous and multiple instances of hybridization among the genetic groups may have occurred during their natural process of differentiation along the Amazon Basin. The gene space of the Iquitos genetic group intersected with a very high intensity (>=92%) with almost all genetic groups, revealing the high diversity of the gene space of this genetic group. Interestingly, intersection between the Criollo gene space and gene space of other genetic groups indicates that Caquetá genetic group encodes the highest percentage (94%) of the genes found in the Criollo genetic group, while the Curaray genetic group, a closely related group of the Criollo group^27^, only covered 88% of the Criollo gene content.

### Functional analysis of variable genes

Gene Ontology (GO) enrichment of protein-coding genes of the variable genome identified significant enriched GO terms (p-value < 0.01) associated to biological processes involved in the plant immunity, gene transcription regulation, plant growth and cellular processes, protein metabolic processes and biosynthesis of aromatic compounds (Table 1) which is consistent with previous findings in other plants (Bayer et al., 2020). Among the most interesting GO terms, 282 variable protein-coding genes were associated with the term "defense response" (GO:006952), suggesting that the variable part of the *T. cacao* genome could play an important role in pathogen resistance processes. Most of them (193) were predicted in the non-reference assembled contigs and were thus not included in the reference genome. We also identified 31 variable genes annotated with "meristem maintenance" (GO:0010073) and 8 genes annotated with "lignin biosynthetic process" (GO:0009809), two key biological processes involved in plant growth and potentially involved in *T. cacao* vigor. Finally, 12 and 11 genes are annotated with "diterpenoid and triterpenoid biosynthetic process" (respectively GO:0016102 and G0:0016104 respectively) indicating that the variable genome may be involved in cocoa flavor compounds.

**Table 1.**
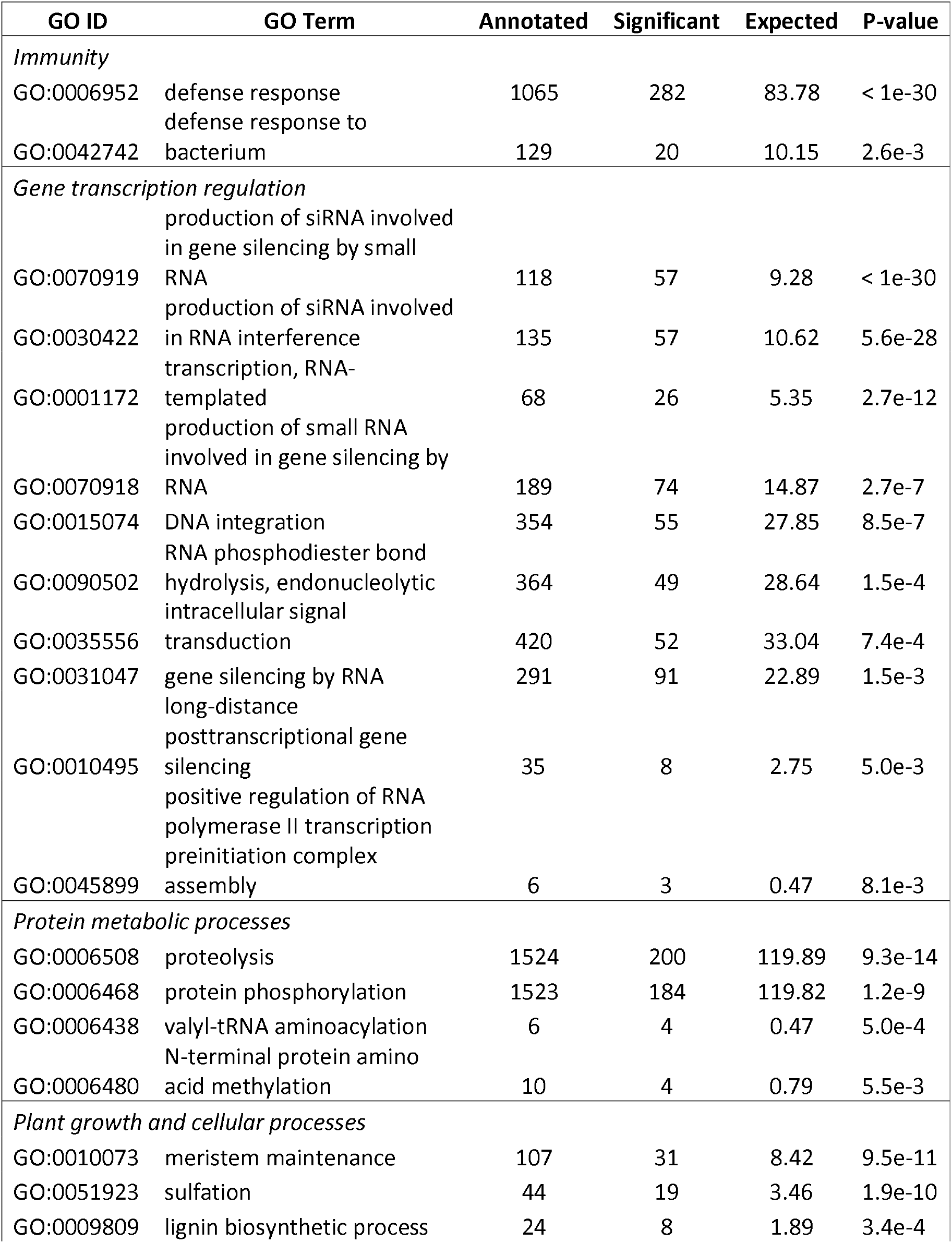
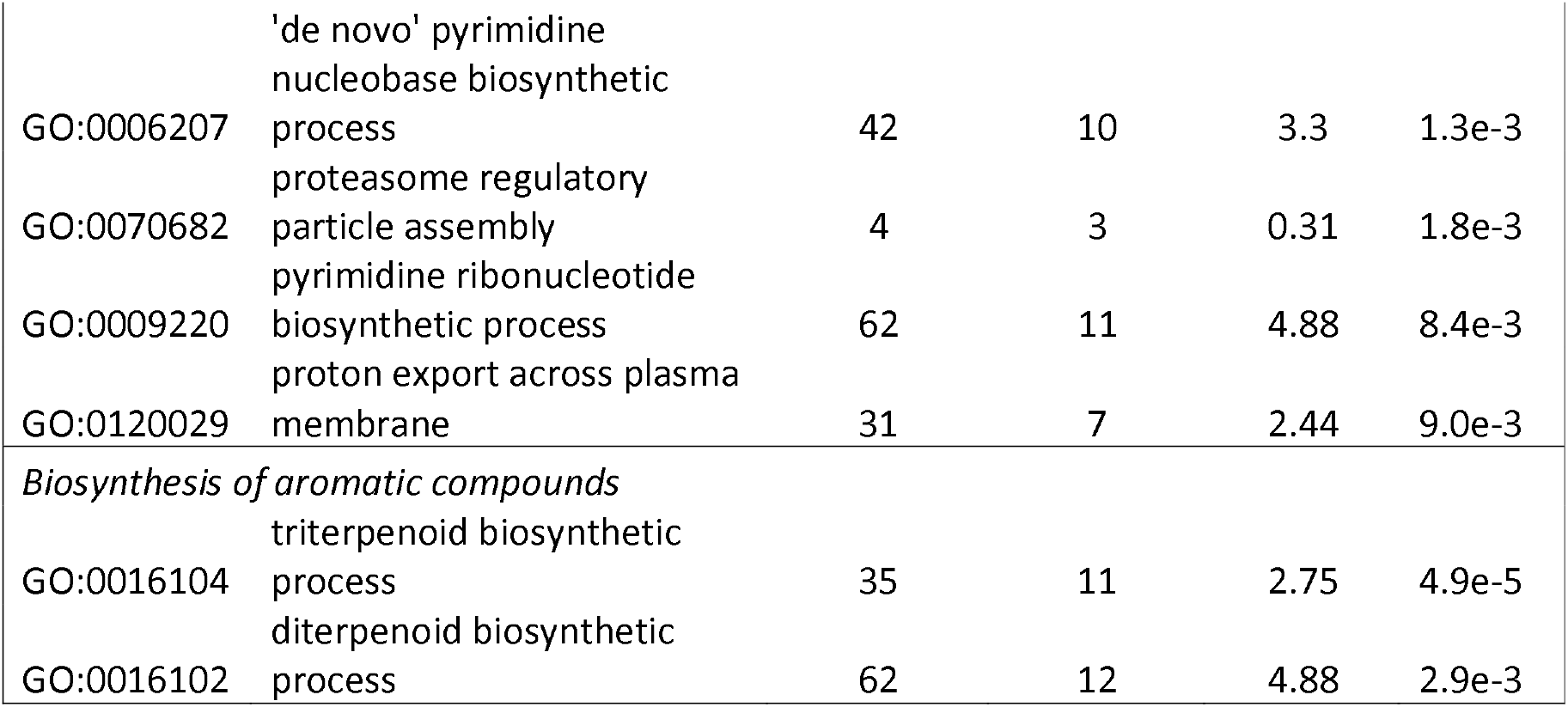
Gene ontology (GO) term enrichment of biological processes found in variable genes.

### Pan-genome-wide analysis of resistance gene analogues (RGAs)

Since the dispensable genome of *T. cacao* seemed particularly enriched in genes involved in plant immunity, we identified at a pangenome-wide scale, RGAs, an important gene family involved in disease resistance in plants. A total of 1385 RGAs were identified in the *T. cacao* pangenome, comprising 654 receptor-like kinase proteins (RLK), 350 nucleotide binding site proteins (NBS), 201 receptor-like proteins (RLP) and 180 transmembrane coiled-coil proteins (TM-CC) (Table 2). Among them, 210 RGAs were variable, indicating that the variable genome may play important roles in determining the level of tolerance to certain diseases observed in particular cocoa accessions. The largest class of variable RGAs corresponded to NBS (30%) followed by RLP (26%) while only 6% and 7% of the RLK and TM-CC resistance gene respectively were found to be variable, indicating that for *T. cacao*, NBS and RLP genes show a greater variability than RLK and TM-CC genes. We then analyzed the distribution of RGAs among the 95 accessions representative of the 15 genetic cluster (with ancestry >0.85). We observed that accessions belonging to the Pangui genetic groups contained the highest number of RGAs (1364), indicating that the Pangui genetic group comprises almost the entire repertoire of the T. cacao RGAs (90% of the T. cacao variable RGAs). This result suggests that germplasm originating from the Pangui genetic group may contain valuable resistance genes that would be worth exploiting. Conversely, 35% of the *T. cacao* variable RGAs were absent in the domesticated Criollo genetic group which comprised the lowest number of RGAs (1312), showing that the domestication process of *T. Cacao* has led to RGA gene loss.

**Table 2:** The number of different RGA candidates and subfamilies found on the T. cacao pangenome and across the 15 genetic groups. The numbers in parentheses represent the number of variable genes. AME=Amelonado, CAQ=Caqueta, CRI=Criollo, CUR=Curaray, APA=Apaporis, GUI=Guiana, IQU=Iquitos, NAP=Napo, MAR=Marañon, NAC=Nacional, NAN=Nanay, PAN=Purus, TIW=Tiwinza

| Individuals | RGA class |  |  |  |  |
| --- | --- | --- | --- | --- | --- |
|  | NBS | RLK | RLP | TM-CC | Total |
| **Pangenome** | 350 (105) | 654 (43) | 201 (52) | 180 (10) | **1385 (210)** |
| **AME** | 321 (76) | 644 (33) | 184 (35) | 178 (8) | **1327 (152)** |
| **APA** | 329 (84) | 648 (37) | 187 (38) | 178 (8) | **1342 (167)** |
| **CAQ** | 337 (92) | 648 (37) | 195 (46) | 177 (7) | **1357 (182)** |
| **CON** | 329 (84) | 647 (36) | 183 (34) | 179 (9) | **1338 (163)** |
| **CRI** | 303 (58) | 649 (38) | 184 (35) | 176 (6) | **1312 (137)** |
| **CUR** | 334 (89) | 649 (38) | 193 (44) | 179 (9) | **1355 (180)** |
| **GUI** | 336 (91) | 648 (37) | 189 (40) | 178 (8) | **1351 (176)** |
| **IQU** | 340 (95) | 650 (39) | 186 (37) | 178 (8) | **1354 (179)** |
| **MAR** | 333 (88) | 648 (37) | 189 (40) | 179 (9) | **1349 (174)** |
| **NAC** | 330 (85) | 652 (41) | 189 (40) | 179 (9) | **1350 (175)** |
| **NAN** | 327 (82) | 646 (35) | 184 (35) | 178 (8) | **1335 (160)** |
| **NAP** | 335 (90) | 650 (39) | 197 (48) | 179 (9) | **1361 (186)** |
| **PAN** | 341 (96) | 651 (40) | 193 (44) | 179 (9) | **1364 (189)** |
| **PUR** | 323 (78) | 648 (37) | 185 (36) | 178 (8) | **1334 (159)** |
| **TIW** | 338 (93) | 649 (38) | 196 (47) | 179 (9) | **1362 (187)** |

### Protein-coding genes under selection during the Criollo domestication

The origin of the domesticated Criollo has been hypothesized to be located in the Upper Amazon region, which includes parts of Ecuador and Colombia^21,28^. To investigate the putative protein-coding genes that might have been under selection during the domestication process of the Criollo group, we compared the gene frequencies between Criollo accessions and accessions native from the putative center of origin of the domesticated Criollo.

We identified in our georeferenced data, a subset of 35 native accessions with Caquetá, Curaray and Napo ancestries, collected by Allen et al. in 1988^29^ in North Ecuador and South Colombia (Figure 4a), hereafter referred to as “native accessions”. Analysis of gene frequencies between accessions from the Criollo group and native accessions (Figure 4b, Supplementary Table 3, Supplementary Table 4) identified 71 genes which present significant higher frequencies in the domesticated Criollo group and therefore would have been positively selected during Criollo domestication and were considered as favorable genes. We also found 624 genes with significantly higher frequencies in native accessions that we regarded as unfavorable or unselected genes during Criollo domestication. No significant enriched Gene Ontology terms was associated with the favorable Criollo genes while defense response was the most enriched term found in the unfavorable genes (Figure 4c). KEGG functional annotation of the unselected genes (Supplementary Table 4) identified 87 genes involved in plant-pathogen interaction pathway (ko04626). Another noteworthy finding was the identification of genes related to the Phenylpropanoid biosynthesis and Flavonoid biosynthesis pathways (ko00940 and ko00941) within the unselected genes. Notably, this included four genes encoding shikimate O-hydroxycinnamoyltransferase (HCT), a central enzyme involved in both pathways. These pathways provide a rich source of metabolites in plants, necessary for the biosynthesis of lignin and are the starting point for the production of many other important compounds, such as anthocyanins^30,31^.

**Figure 4.**
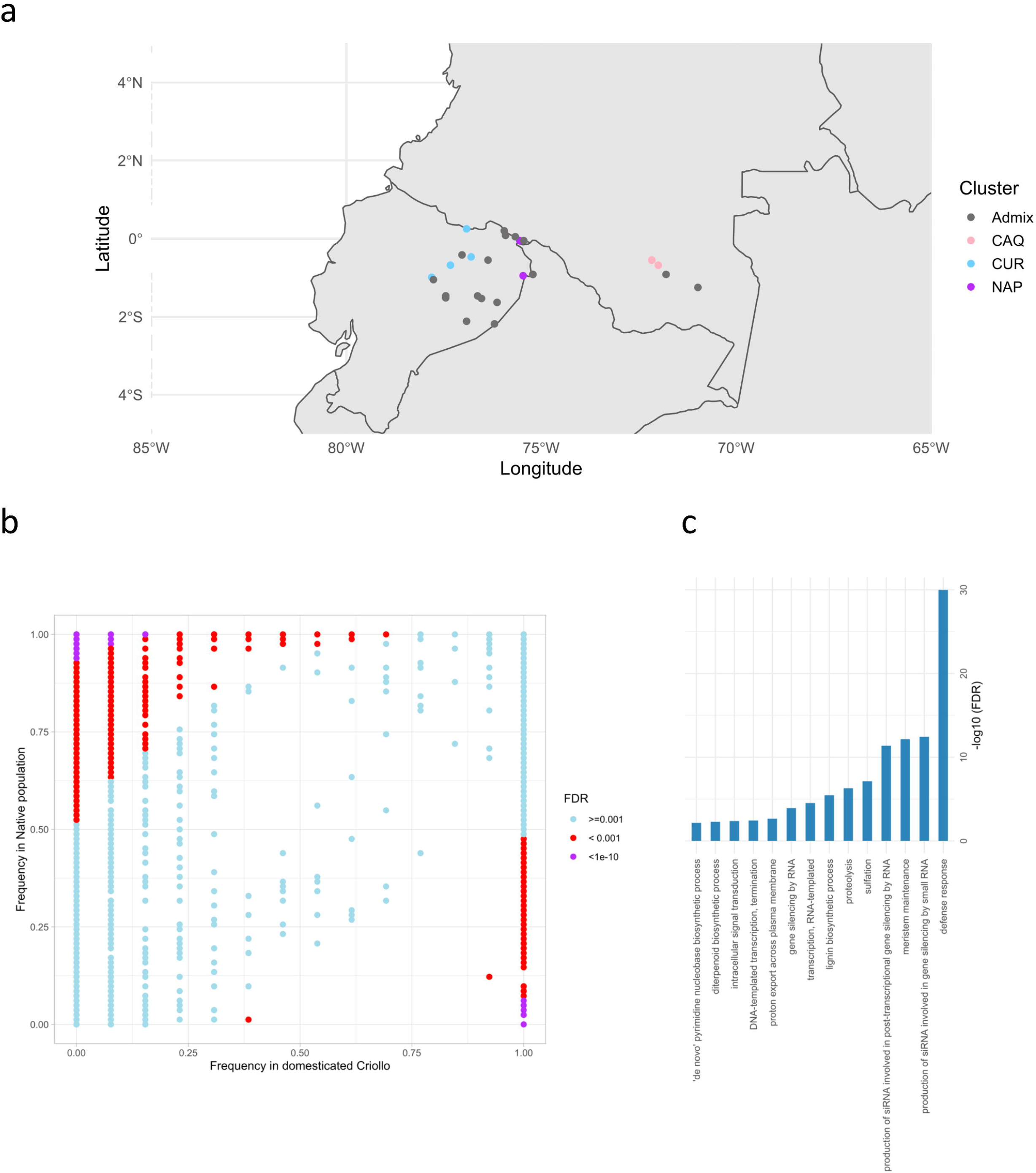
Protein-coding genes under selection during the Criollo domestication. (a) Geographic location of the 35 *T. cacao* accessions native from the putative center of origin of the domesticated Criollo. (b) Scatter plot showing gene occurrence frequencies in the population of 35 accessions native from the putative center of origin of the domesticated Criollo and in the 13 accessions of the Criollo group. (c) Enriched GO terms in unfavorable genes during Criollo domestication.

## Discussion

In this study, we analyzed the genetic diversity of the *Theobroma cacao* species through the prism of its pangenome, define here as all the genes contained in the species, whether present in all individuals (core genome) or absent in one or more individuals (variable genome). For this purpose, we analyzed a population of 216 cacao trees covering a wide genetic diversity and including, for the majority of the studied accessions, subspontaneous genetic material, collected during surveys in the main geographical areas of origin of *Theobroma cacao*^21,29,32–34^. The construction of the pangenome has increased the genome of the *Theobroma cacao* species, with 48.2 Mb of new genomic sequences and 1,407 new candidate protein-coding genes absent from the B97-61/B2 reference genome^9^. Analysis of the cocoa pangenome composition showed, unusually, a very high content of core genes (90.8%), in contrast to what has been observed in other plant species such as melon^35^ (73.5%), white lupin^36^ (78.5%), tomato^37^ (74.2%) and cotton^38^ (74.1%). We also highlighted that very few genes were specific to the genetic groups identified and that none of the genes were specific to a single accession. It could therefore be hypothesized that the evolutionary processes behind the diversification of the cocoa tree through genetic drift or domestication have had little impact on the gene content between accessions, and that the species has not yet suffered any major gene loss.

Recent research^39^ has also suggested that genetic mixing among various *T. cacao* genetic groups might have occurred several millennia ago due to human influence. This gene flow between ancestral genetic groups would thus have limited the effects of their original differentiation and led to today’s *T. cacao* populations. Clement et al. (2015)^40^ proposed that Amazonia played a pivotal role as a major world center for plant domestication. Human societies in the Amazonian regions have been significantly altering species composition in various ecosystems since the Pleistocene to the Early Holocene, simultaneously amassing valuable crop genetic resources as their societies expanded. Intensive trading interactions, including the exchange of plants, facilitated by the extensive river networks across Amazonia, occurred between distant human populations thus favorizing gene flow between contrasted genetic groups. For instance, an important center of plant genetic resources was identified in the Iquitos region of Peru^40^, where multiple *T. cacao* genetic groups have been documented^22^. The *T. cacao* Iquitos genetic group, characterized by its exceptionally diverse gene content that extensively overlaps with nearly all other genetic groups, may have originated through the hybridization of multiple ancestral genetic groups. This hypothesis is reinforced by the high rate of heterozygosity found in Iquitos accessions (49.3%), as reported by Fouet et al. (2022)^21^.

Another noteworthy finding related to the Guiana genetic group, which, despite its remote location in the far eastern region of the Amazon basin (French Guiana), shares a close genetic relationship with the Peruvian Nanay, Marañon, Amelonado and Iquitos accessions, the latest comprising 95% of the gene space of the Guiana group. The Guiana genetic group is also one of the group with the fewest gene content and was found to be highly homozygous (Observed Heterozygosity of 7.9%)^21^. Archaeological research evidenced the presence of human populations settle in French Guiana dating from 4500 to 2000 BC^41^. The distribution of linguistic groups in contact in Amazonia^42^ shows that the Tupi language family extended from Iquitos until to the mouth of the Amazon, and on either side including French Guiana, thus supporting possible contact between these two regions. All these results suggest that accessions from the Iquitos region could have been transported by humans to French Guiana through Amazonian rivers, initiating a domestication process that may have involved gene losses.

### Evolutionary origin of the Criollo genomes and Impact on gene content

Examination of the distribution of gene PAVs among the 15 genetic groups revealed that accessions belonging to the early cultivated Criollo genetic group exhibited the most restricted gene catalogs (29,298). Gene losses have been particularly observed in plants and animals in populations subjected to domestication or genetic improvement^37,38,43,44^. The gene losses observed in the Criollo genetic group thus provide further evidence of domestication processes that led to its genetic differentiation, such as those already reported in previous studies with SSR and SNP markers^27,28^. While there is a scientific consensus suggesting that Criollo was domesticated from cacao populations originating in the Upper Amazon^21,27,28^, the genetic origin of the Criollo group remains controversial. Some authors^20,27^ argued that Criollo could have been domesticated from ancient Curaray germplasm, a genetic group described from North Ecuador to South Colombia. A recent study based on molecular analysis of cocoa trees native to the upper Amazon suggests that Criollo is most closely related to the Caqueta genetic material, collected in southern Colombia^21,29^. Using PAVs, the results presented in our study confirmed the closer genetic similarity of Criollo accessions to accessions from the Caqueta group. We found that the Caqueta group contains 94% of the Criollo gene space, in contrast to the Curaray germplasm, which encodes only 88% of the genes found in Criollo. This greatest similarity is also reinforced in the Neighbor-Joining analysis where Criollo accessions are closely related to Caqueta. All these results suggest that the domestication of Criollo could have been initiated from individuals close to the Caqueta genetic group.

We further investigated putative protein-coding genes under selection during the domestication process of the Criollo group by identifying significant different frequencies between accessions from the Criollo group and subspontaneous accessions collected in the putative area of origin of the Criollo group. We identified 71 positively selected genes and 624 negatively selected genes during Criollo domestication. The domestication process led to a significant loss of genes involved in defense response, which could partially explain the high disease susceptibility of Criollo observed in the fields^45,46^. Nowadays, Criollo cocoa trees are grown in very restricted areas and has been gradually replaced by Criollo hybrids (Trinitario) which provide more disease resistance and productivity^23,28,47^.

The analysis of the unselected genes during Criollo domestication also revealed the negative selection of genes involved in the metabolic pathways of phenolic compounds, such as Phenylpropanoid and Flavonoid biosynthesis pathways. The flavonoids constitute the largest and most diverse group of phenolic compounds in cocoa beans, among which anthocyanins, responsible for the purple color of cacao beans, are generally present in large quantities^48^. One of the main desirable criteria of the Criollo varieties is their beans specificities, which are white or very slightly colored and recognize for their high-quality flavor properties with low astringency and low bitterness due to low polyphenolic compounds content such as anthocyanins^49^. These particular features of the Criollo beans let us hypothesized that early selection of Criollo varieties during domestication has been done on their bean’s qualities. Furthermore, several authors have observed that Criollo exhibits very low vigor and is particularly challenging to propagate vegetatively^23,47,50,51^. As phenolic compounds are also involved in lignin production processes and in regulating cell growth, the selection of Criollo on the criterion of white bean color and low bitterness might also have been associated with a counter-selection of genes involved in plant vigor.

### Potential impact of PAVs on disease resistance breeding

Pest and disease resistance is one of the major targets of most cocoa breeding programs^52^. To investigate the resistance gene resources available in *T. cacao*, we used the pan-genome to identify resistance gene analogues (RGAs) and analyzed their distributions and PAVs among the different genetic groups. We found that a relatively substantial fraction of resistance genes resides in dispensable loci (15%), underscoring the importance of conserving and utilizing *T. cacao* genetic diversity for disease control.

The most common sources of disease resistance used presently by breeding programs belong to accessions from Marañon, Nanay, Contamana and Iquitos ancestries^3^. These accessions were collected in Peru by Pound in 1938 and 1943^33,53^ to search for *T. cacao* accessions resistant to witches’ broom disease. This collected genetic material also had the advantage of being resistant or tolerant to other widespread diseases, such as black pod disease caused by several *Phytophthora* species. Other sources of resistance to *Phytophthora* were more recently observed in subspontaneous *T. cacao* accessions from Guiana ancestry^54–56^. Surprisingly, no significant expansion of RGAs was found in accessions from these genetic groups. Interestingly we found that accessions from Pangui and Tiwinza ancestries, collected since 2010 in the Ecuadorian provinces of Zamora-Chinchipe, Morona- Santiago, and Pastaza^21^ contained the largest number of RGAs. This result suggests that new sources of disease resistance could be available in this germplasm, currently being evaluated at INIAP research stations in Ecuador.

## Materials and methods

### Plant material

In this study, leaf tissues were collected leaf from 185 accessions maximizing the available of *Theobroma cacao* as described by Fouet *et al.* (2022)^21^, and from 3 accessions from *Theobroma cacao* related species, *Theobroma bicolor*, *Theobroma grandiflorum* and *Theobroma speciosum,* used as outgroups (Supplementary Table 1).

### Genome sequencing and reads cleaning

Genomic DNA was extracted and purified for each accession using the protocol published by Risterucci *et al.* (2000)^57^ and modify by Allegre *et al. (* 2012)^58^. Sequencing libraries were prepared as previously described in Alberti *et al.* (2017)^59^ using an ‘on beads’ protocol. Each library was sequenced using 151 base-length read chemistry on a paired-end flow cell on the Illumina NovaSeq6000 (Illumina, USA). After Illumina sequencing, an in-house quality control process was applied to the reads that passed Illumina quality filters. The first step discarded low quality nucleotides (Q < 20) from both ends of the reads. Next, Illumina sequencing adapters and primer sequences were removed from the reads. Then, reads shorter than 30 nucleotides after trimming were discarded. Trimming and removal steps were achieved using custom-developed software based on the FastX package (https://github.com/institut-de-genomique/fastxtend). Finally, using SOAP aligner^60^, read pairs that aligned to the phage phiX genome and the Enterobacteria phage PhiX174 reference sequence (GenBank: NC_001422.1) were identified and discarded.

To complement the genetic diversity of the dataset, Illumina genome sequencing data (150bp paired end reads) of 31 accessions^24^ were retrieved from NCBI SRA database (BioProject PRJNA558793). Quality control of sequencing reads was performed using the same protocol mentioned above, and sequencing depth of these accessions was normalized to 64X with fastq-sample (https://github.com/fplaza/fastq-sample), corresponding to the third quartile of the sequencing coverage of the newly sequenced 185 *T. cacao* accessions.

### Pangenome assembly and annotation

We applied an assemble-then-map approach. For each accession, cleaned Illumina reads were *de novo* assembled using Megahit v1.2.9^61^ with a minimum length of contigs of 500bp (option --min-contig-len=500). The completeness and accuracy of the assembly were assessed using BUSCO v5.0.0^62^ in genome mode with the lineage dataset eudicots_odb10.

Contigs were then aligned to a reference dataset comprising the "B97-61/B2" reference genome^9^ and the complete *Theobroma cacao* chloroplast genome (GenBank: HQ244500.2) using NCBI Megablast v2.9.0^63^ with e-value threshold of 1e-5. Sequences ≥ 500bp and below 90% of identity with alignment with the reference dataset were considered novel (non-reference) and retained for further analysis. The non-reference sequences of each genotype were then searched against the GenBank nucleotide collection NT using Megablast (parameter -evalue=1e-5). Subsequently, we used the BASTA taxonomic classification tool^64^ with default parameters was employed to identify and eliminate sequences not belonging to the green plants clade or those related to known plant mitochondrial or chloroplast genomes. The cleaned nonreference sequences of all accessions were then combined to reduce sequence redundancy with cd-hit-est of CD-HIT package v4.8.1^65^ using a sequence identity threshold of 90%. The nonreferenced, cleaned and nonredundant dataset was then combined to the "B97-61/B2" reference genome to form the *Theobroma cacao* pan-genome.

Automatic gene prediction was performed on the pan-genome with the integrative EuGene Eukaryotic Pipeline (EGNEP) v1.5.1 including EuGene v4.2a^66^. A custom statistical model for splice-site detection was applied to improve the quality of prediction based on the published reference genome^9^. Three protein databases were aligned to contribute to translated regions detection, (i) the proteome of *T. cacao* "B97-61/B2" reference genome, (ii), UniProt/SwissProt Viridiplantae, (iii) the proteome of *T. cacao* cultivar Matina 1-6^10^. Two transcriptome databases were used for transcriptional evidence, one representing the transcriptome of the "B97-61/B2" reference genome^8^ and the second composed of Expressed Sequence Tags (ESTs) derived from diverse genotypes, tissues and experimental conditions^67^. Proteins hits from Repbase were filtered prior to functional annotation. Gene functions were assigned through InterProScan domain searches and similarity searches against the UniProt/Swissprot and UniProt/TrEMBL databases (BlastP). Gene Ontology (GO) annotation of the pan-genome protein-coding genes was implemented using OmicsBox/Blast2GO v2.1^68^. Prediction of plant resistance gene analogs was carried out with the RGAugury pipeline^69^.

### Analysis of Gene Presence/Absence Variation (PAV)

Sequencing reads from each accession were aligned to the *T. cacao* pan-genome using Bowtie2^70^ v2.4.2 under the “very-sensitive-local” mode. Gene PAV was determined based on the cumulative coverage of exons of each protein-coding gene model in each accession using SGSGeneLoss^71^ v0.1. A gene was considered present if a minimum of 20% of its exons were covered by at least two reads (options minCov = 2, lostCutoff = 0.2), otherwise it was considered absent.

Estimation of ancestral populations and ancestry coefficients was calculated from the PAV matrix of the 216 *Theobroma cacao* accessions using the admixture analysis provided by the LEA R package^72^ v3.8.0. The sparsen nonnegative matrix factorization (snmf) function was first executed with K value ranging from 2 to 30 with 100 repetitions. Then, the lowest value of the cross-entropy criterion found between the different K values was used to select the number of ancestral populations that best explains the genotypic data. Accessions having ancestry proportion > 85% were considered representative of an ancestral population, otherwise were considered as admixed.

The population structure was then analyzed with principal component analysis (PCA) implemented in the LEA package. The number of significant components was computed with the Tracy-Widom test (pvalue<0.01).

Neighbor-Joining phylogenic analysis was conducted using Darwin Software^73^ v6.0.21. The distance matrix was calculated from the PAV matrix of the 216 *T. cacao* accessions and 3 accessions from outgroups (*T. bicolor*, *T. grandiflorum* and *T. speciosum*). Distances were computed using the Sokal & Michener (simple matching) algorithm as implemented in Darwin software. The first topology was calculated using 95 accessions exhibiting ancestry > 0.85 with the weighted Neighbor-Joining method and 1000 bootstrap. The resulting NJ tree was then used as constraints to build the phylogenetic tree of the remaining 121 *T. cacao* accessions using the “Neighbor-Joining under topological constraints” method implemented in Darwin software. Finally, the 3 outgroup accessions were grafted a posteriori on the resulting NJ tree to avoid disturbance of these distant accessions on the resulting tree.

The identification of gene PAVs under selection during the process of domestication of the Criollo genetic group was done according to the methodology described in Gao et al., (2019)^37^. Basically, the frequency of each gene in the Criollo population and in the native population from the putative center of origin of the domesticated Criollo were first computed. Then, Fisher’s exact test was applied to test the significance of the difference in gene frequency between each group and p-values were adjusted using Benjamini and Hochberg False Discovery Rate (FDR) method. Genes exhibiting a FDR less than 0.001 and a frequency fold change exceeding 2 were considered to be at significantly different frequencies between the two groups and were identified either as favorable (selected in the Criollo population) or unfavorable (unselected) genes. GO enrichment was performed using TopGO R package v.2.48.0 with the Fisher’s exact statistical test and the “elim” method. KEGG annotation was computed by eggNOG-mapper^74^ v2.1.9.

## Availability of data and material

Raw sequence reads were deposited in the Sequence Read Archive (SRA) of the National Center for Biotechnology Information (NCBI) (BioProject: PRJNA880462). Pangenome assembly, Gene structure and functional and additional data sets are made available on Download section of the Cacao Genome Hub (https://cocoa-genome-hub.southgreen.fr/download).

## Authors’ contributions

X.A. and G.D performed bioinformatic and statistical analysis. O.F and K.L. performed wet-lab experiment and sequencing. O.F. and G.R.L. contributed to sample collection. M.R. and B.R. assisted with data analysis and manuscript revision. X.A. and C.L. conceived research designed analysis and wrote manuscript.

## Acknowledgements

This work has been realized with the support of MESO@LR-Platform at the University of Montpellier, the financial support of CIRAD, and the technical support of the bioinformatics group of the UMR AGAP Institute, member of the French Institute of Bioinformatics (IFB) – South Green Bioinformatics Platform.

We thank CRC (Cocoa Research Center – Trinidad and Tobago), CATIE (Centro Agronómico Tropical de Investigación y Enseñanza-Costa Rica) and INIAP (Instituto Nacional de Investigaciones Agropecuarias – Ecuador) for providing leaf samples of *T. cacao* accessions studied in this work.

We thank Kelly Colonges for her help in DNA extraction.

We thank the I-Site MUSE and Valrhona for their financial support of this project. This work, part of the MUSE Amazcacao project, was publicly funded through ANR (the French National Research Agency) under the "Investissement d’avenir" program with the reference ANR-16-IDEX-0006.

---

**Supplementary Figure 1**

Word clouds of Gene ontology terms found in the non-reference genes belonging to different biological processes.

**Supplementary Figure 2**

Model-based clustering (from K=2 to K=15) of the 216 *T. cacao* accessions based on the 2,802 gene PAVs.

**Supplementary Figure 3**

Principal component analysis of the 216 *T. cacao* accessions based on the 2,802 gene PAVs.

