## Supplementary figures for "Pangenomic exploration of *Theobroma cacao*: New Insights into Gene Content Diversity and Selection During Domestication"

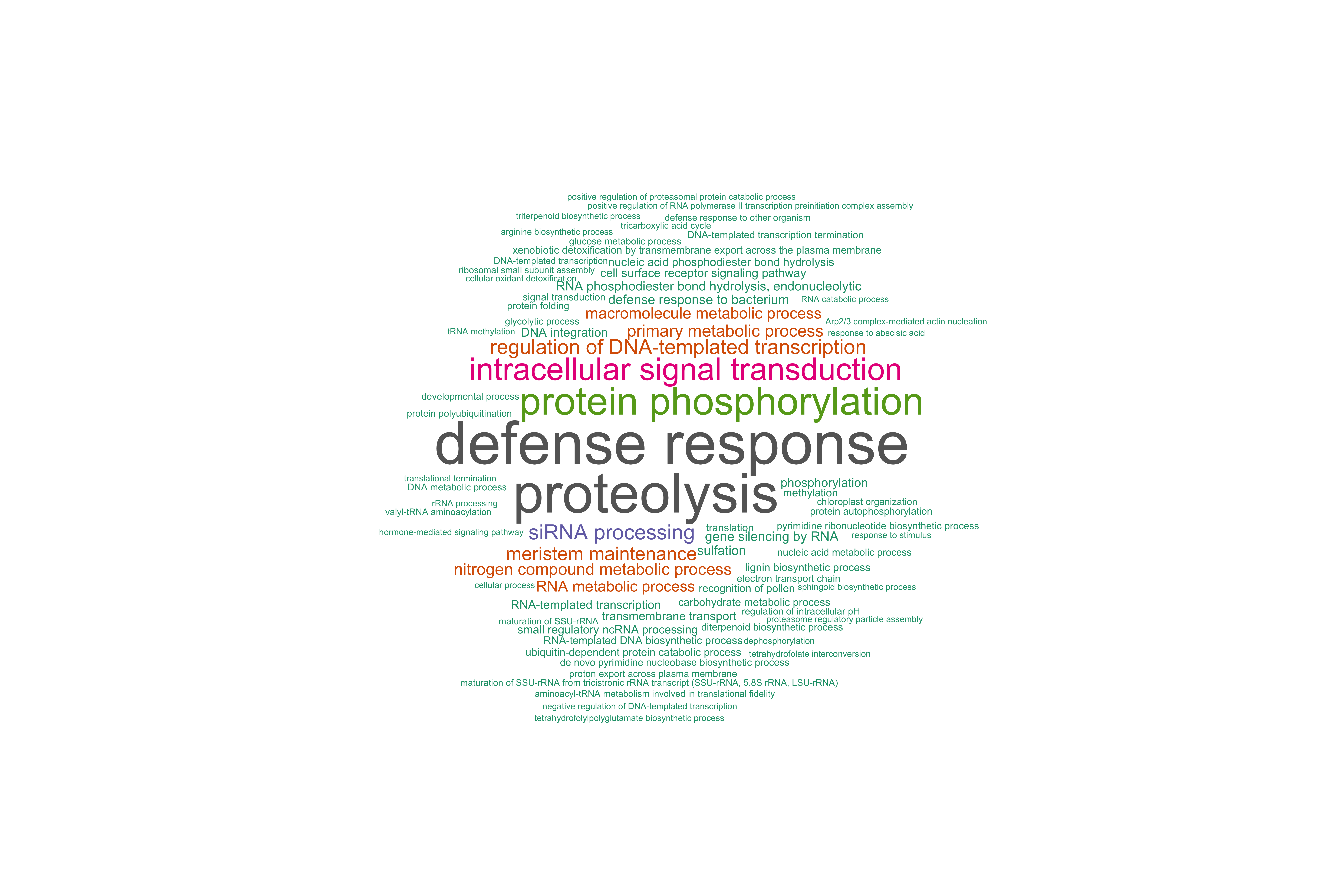

**Supplementary Figure 1**

Word clouds of Gene ontology terms found in the non-reference genes belonging to different biological processes.

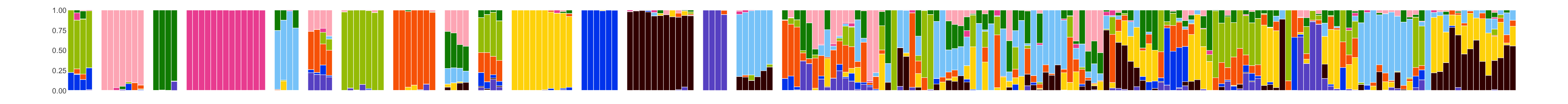

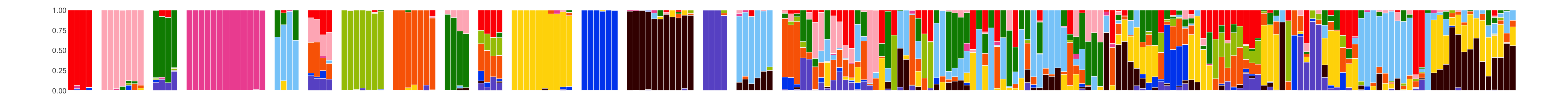

K=15

K=14

K=13

K=11

K=10

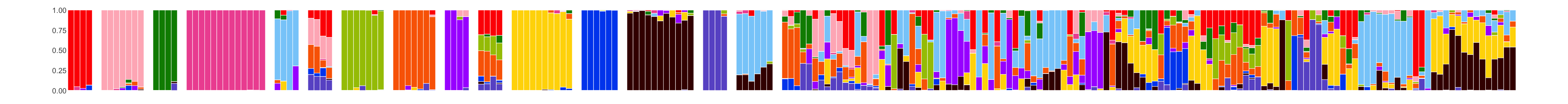

K=12

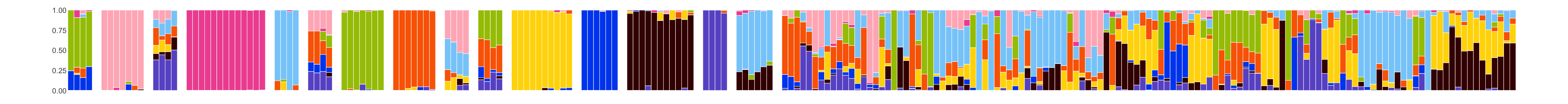

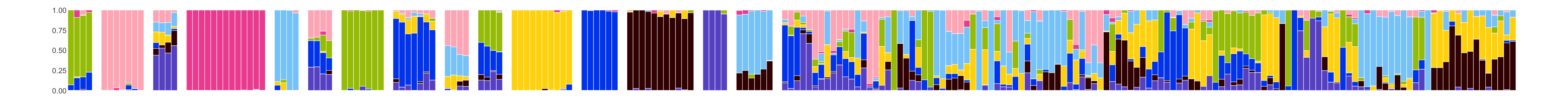

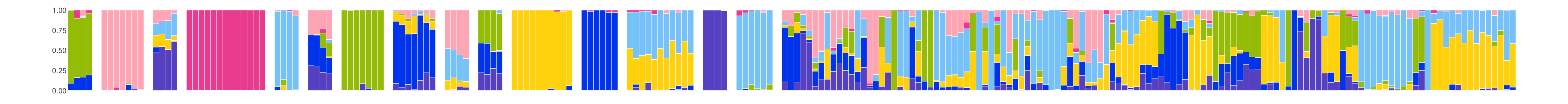

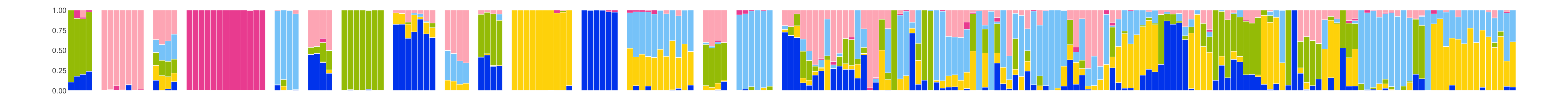

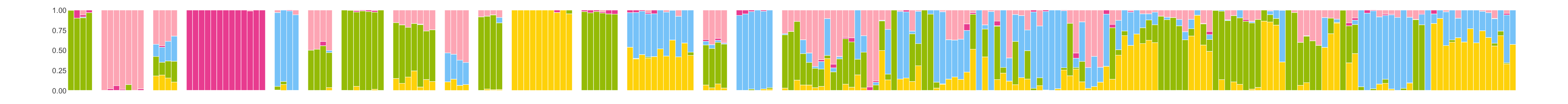

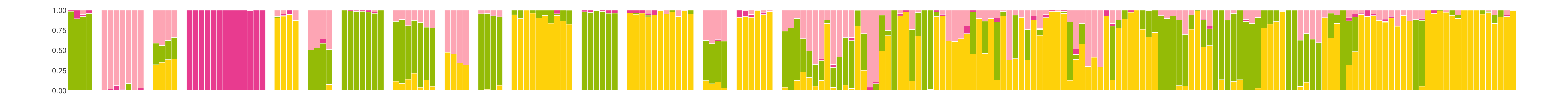

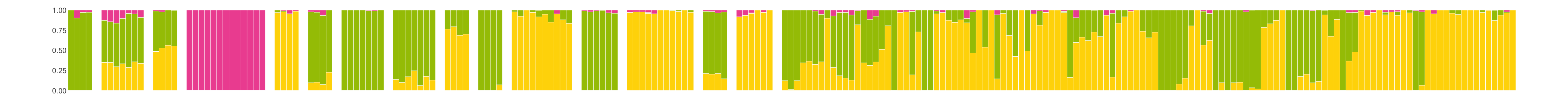

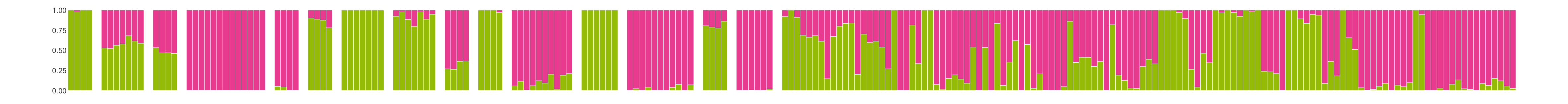

K=9

K=8

K=7

K=6

K=5

K=4

K=3

K=2

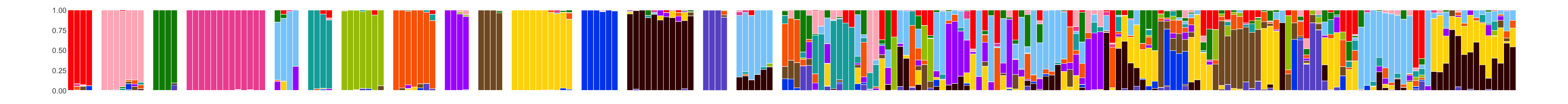

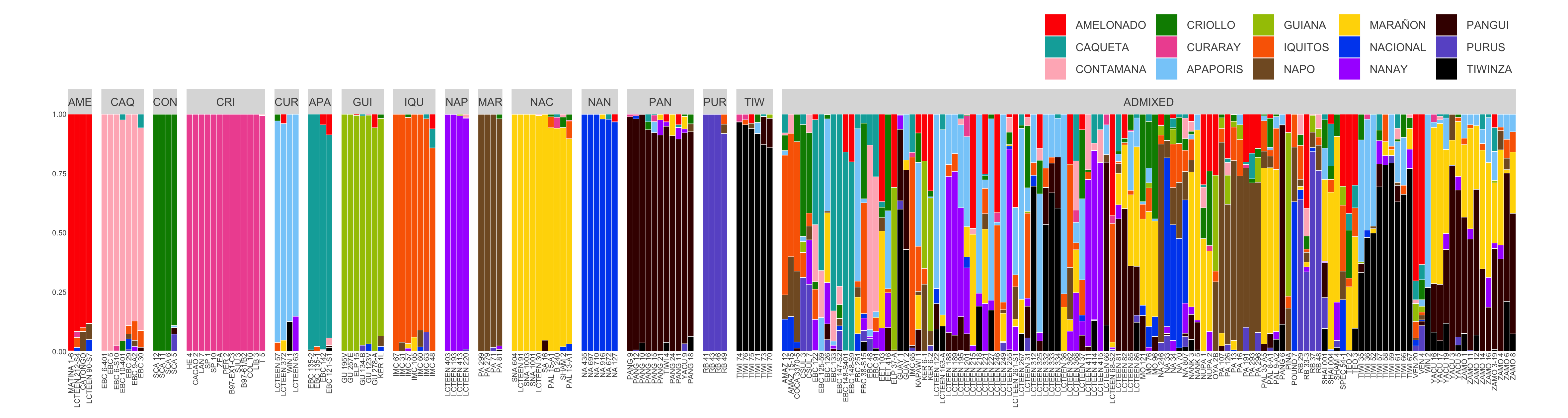

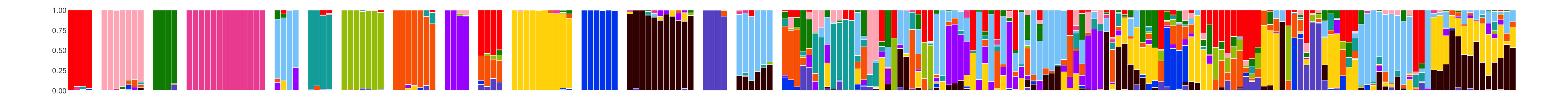

**Supplementary Figure 2**

Model-based clustering (from K=2 to K=15) of the 216 *T. cacao* accessions based on the 2,802 gene PAVs.

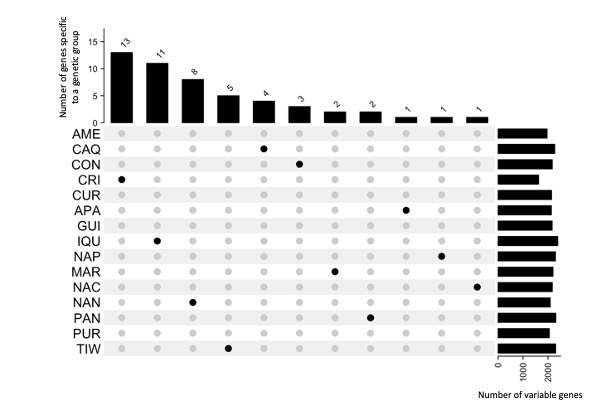

**Supplementary Figure 3**

UpSet plot displaying the number of group-specific genes for the 15 genetic groups. Amelonado, Curaray, Guiana and Purus genetic groups have no group-specific genes.
